# Pupil Size Reflects Exploration and Exploitation in Visual Search (and It’s Like Object-Based Attention)

**DOI:** 10.1101/2021.02.05.429946

**Authors:** Franziska Regnath, Sebastiaan Mathôt

**Affiliations:** University of Amsterdam; Department of Experimental Psychology, University of Groningen

**Keywords:** locus coeruleus, pupil size, adaptive gain theory, visual search, eye movements

## Abstract

The adaptive gain theory (AGT) posits that activity in the locus coeruleus (LC) is linked to two behavioral modes: exploitation, characterized by focused attention on a single task; and exploration, characterized by a lack of focused attention and frequent switching between tasks. Furthermore, pupil size correlates with LC activity, such that large pupils indicate increased LC firing, and by extension also exploration behavior. Most evidence for this correlation in humans comes from complex behavior in game-like tasks. However, predictions of the AGT naturally extend to a very basic form of behavior: eye movements. To test this, we used a visual-search task. Participants searched for a target among many distractors, while we measured their pupil diameter and eye movements. The display was divided into four randomly generated regions of different colors. Although these regions were irrelevant to the task, participants were sensitive to their boundaries, and dwelled within regions for longer than expected by chance. Crucially, pupil size increased before eye movements that carried gaze from one region to another. We propose that eye movements that stay within regions (or objects) correspond to exploitation behavior, whereas eye movements that switch between regions (or objects) correspond to exploration behavior.

**Public Significance Statement:** When people experience increased arousal, their pupils dilate. The adaptive-gain theory proposes that pupil size reflects neural activity in the locus coeruleus (LC), which in turn is associated with two behavioral modes: a vigilant, distractible mode (“exploration”), and a calm, focused mode (“exploitation”). During exploration, pupils are larger and LC activity is higher than during exploitation. Here we show that the predictions of this theory generalize to eye movements: smaller pupils coincide with eye movements indicative of exploitation, while pupils slightly dilate just before make eye movements that are indicative of exploration.

## Pupil Size Reflects Exploration and Exploitation in Visual Search (and It’s Like Object-Based Attention)

The eyes’ pupils become smaller (constrict) in response to brightness or when focusing on a nearby object. Conversely, pupils become larger (dilate) in darkness or when shifting gaze to a far-away object (for a review, see Mathôt, 2018). In addition, pupil dilation is linked to increased autonomic arousal. For instance, Bradley et al. (2008) showed that participants’ pupils dilated more while viewing pictures that were either pleasant (e.g., rafters) or unpleasant (e.g., a burn victim) compared to neutral (e.g., a man). Importantly, these changes in pupil diameter were related to participants’ increased arousal in response to the emotional pictures, and were independent of the pictures’ valence (positive versus negative). In addition, pupil dilation covaried with concurrently measured skin-conductance, which suggests that arousal-related pupil responses are predominantly mediated by increased activity in the sympathetic nervous system.

### The Adaptive Gain Theory: A Link Between Locus Coeruleus Activity, Pupil Size, and Behavior

Current theories suggest that arousal-related changes in pupil diameter are closely linked to activity in the locus coeruleus-norepinephrine (LC-NE) system (Murphy et al., 2014). The LC is a brain-stem nucleus containing noradrenergic neurons that send widespread projections throughout the central nervous system. As such, the LC appears to play a central role in the regulation of various physiological and psychological processes, including general arousal and attention control, the sleep-wake cycle, stress responses, and emotional states (for reviews see, Aston-Jones & Cohen, 2005; Sved et al., 2002; Zitnik, 2016). Because of the correlation between LC activity and pupil size, pupillometry is frequently used to indirectly track activity of the LC-NE system.

The adaptive-gain theory (AGT), first proposed by Aston-Jones and Cohen (2005), provides a more refined framework to understand the three-way link between the LC-NE system, pupil size, and behavior; specifically, the AGT posits a distinction between two modes of behavior: exploitation and exploration. Exploitation is characterized by a narrow focus of attention on the current task, and is accompanied by intermediate tonic (sustained) but high phasic (bursty) LC activity, and also by intermediate size pupils that dilate strongly in response to stimuli. In contrast, exploration is characterized by distractibility and a tendency to switch between different tasks, and is accompanied by high tonic and low phasic LC activity, and also by large pupils that dilate less strongly in response to stimuli.

The alternation between exploitation and exploration modes of behavior is thought to be driven by changes in task utility. That is, as a task becomes less rewarding, we tend to disengage from it and to try out other tasks that may offer higher rewards. Therefore, switches between exploitation and exploration could offer adaptive value by promoting withdrawal from fruitless tasks in favor of more rewarding tasks (Aston-Jones & Cohen, 2005; Aston-Jones et al., 1999; Jepma & Nieuwenhuis, 2011).

Pajkossy et al. (2017) tested the AGT with a computerized version of the Wisconsin Card Sorting Task, in which participants were presented with four cards, each portraying symbols differing in color, shape, and number. Based on a hidden rule, participants had to match a fifth card to one of the four cards (e.g., three green triangles matched with three red circles, the shared dimension being the number of symbols). However, the matching rule shifted after a certain number of trials, and subjects had to discover the new rule through trial-and-error. Thus, each rule change was accompanied by a loss of reward (i.e., no positive feedback) due to a mismatch of the cards, thereby triggering a switch from exploitation (sticking to one matching rule) to exploration behavior (trying out new matching rules). That is, during exploration (in this task), participants try out other cards by matching symbols based on different dimensions until the new rule is identified, and behavior shifts back to exploitation. Consistent with the AGT, this study showed that pupil size was larger during periods of exploration behavior than during periods of exploitation behavior.

The relationship between LC-NE activity, pupil size, and behavior is largely based on correlational studies such as this one (Pajkossy et al., 2017). However, there is some evidence for a causal link from research using animal models (for a review, see Waterhouse & Navarra, 2019). For instance, Kane et al. (2017) used designer receptors to directly increase tonic firing of neurons in the LC-NE system of rats while the animals completed a decision-making task. In line with the AGT, rats exhibited diminished exploitative behavior during stimulated LC tonic activity, as reflected by worsened performance during the task and early withdrawal from a currently rewarding source in order to pursue an alternative, unknown reward at a different location. That is, there is limited evidence to suggest that LC-NE activity causes, rather than merely correlates with, exploitation or exploration modes of behavior.

### Cognitively Driven Pupil Size Changes as Sensory Tuning

The AGT provides a compelling description of how pupil size correlates with LC-NE activity and behavior. However, the theory does not speak to the question of why there is such a correlation. Is there a functional reason for why a switch to an exploration mode of behavior is accompanied by pupil dilation?

Our visual system balances sharpness of vision (i.e., visual acuity) and detection of faint stimuli (i.e., visual sensitivity) to achieve optimal vision in a given situation. Specifically, small pupils provide high visual acuity (by reducing optical distortions of the lens), and thus facilitate stimulus discrimination. In contrast, large pupils provide high visual sensitivity (by allowing more light to enter the eye) and thus facilitate stimulus detection, which is especially beneficial in darkness (e.g., see Mathôt & Ivanov, 2019; Woodhouse, 1975). A fully dilated pupil can constrict from about 8 mm to approximately 2 mm when exposed to intense brightness (Walker et al., 1990) and this sixteenfold decrease in pupil area has a substantial impact on vision. However, cognitively driven changes in pupil size are far smaller (±0.5 mm). Some authors have even proposed that these pupil-size changes are too small to be functional, and must therefore be a non-functional by-product of other nervous system processes (e.g., Beatty & Lucero-Wagoner, 2000). However, other authors have suggested that even tiny changes in pupil size could serve to optimize vision, albeit in more subtle ways (e.g., Ebitz & Moore, 2019; Mathôt, 2020).

Mathôt (2020) proposed that cognitively driven changes in pupil diameter could reflect a form of “sensory tuning”: a subtle adjustment of the eyes to optimize their properties for the situation. That is, a calm focus on central vision during exploitation favors visual acuity, while a more distractible or vigilant state during exploration would coincide with a shift of focus to the periphery and an enhancement of visual sensitivity. Such a distinction between central and peripheral focus is in line with findings by Daniels et al. (2012), who demonstrated that pupil size differed for narrowly focused versus broadly spread attention. More specifically, they found that participants’ pupils were relatively constricted when attention was narrowly allocated to central elements on the screen, while their pupils dilated when participants focused their attention more broadly to elements located in the periphery.

Depending on the situational demands, even a slight improvement in vision could be the deciding factor in an extreme situation (e.g., spotting the presence of a predator). Therefore, it is plausible that even a minor advantage could have been valuable at some point in evolutionary history. For instance, Charles Darwin (1859) highlighted that “[…] any being, if it vary however slightly in any manner profitable to itself, under the complex and sometimes varying conditions of life, will have a better chance of surviving, and thus be naturally selected” (p. 5).

In summary, the link between behavioral modes and pupil size could reflect a subtle optimization of vision (Mathôt, 2020). If so, differences between exploitation and exploration modes of behavior should also be evident in a visual-search task, which is a highly visual task.

### The Present Study

The goal of the present study was to determine whether changes in eye-movement behavior and pupil size could be captured by predictions of the AGT. We used a visual-search task in which participants searched for a single target letter among 224 distractor letters. The target and distractors were small and numerous enough so that participants had to make many eye movements across the search display in order to find the target. Importantly, we divided the search display into four differently colored regions, which randomly changed in shape and size after each trial. Although these regions were not relevant for successful task completion, we expected that viewing behavior would still be guided by these regions.

Therefore, we hypothesized that participants would stay within regions for longer than would be expected by chance. Following the AGT, we also expected pupils to dilate just before participants switched from one region to another, reflecting a shift from exploitation to exploration. That is, in this visual-search task, eye movements within a region would represent exploitation (i.e., a calm, narrow focus on letters within the same region), whereas eye movements between regions would reflect exploration (i.e., a broad focus on stimuli outside the attended region). In sum, we predicted that within-region eye movements are a form of “micro-exploitation” that should be accompanied by smaller pupils. In contrast, eye movements between regions would be a form of “micro-exploration” that should be accompanied by larger pupils.

## Method

### Participants

Thirty-six undergraduate psychology students (24 women, 12 men) from the University of Groningen participated in the experiment in exchange for course credits. Only subjects with normal or corrected-to-normal vision were invited to take part in the study. Prior to participation, all subjects received research information and provided written informed consent. The experiment was approved by the Ethical Committee of the department of Psychology at the University of Groningen (approval code 18286-S Tetris). Since we did not have any a-priori estimate of expected effect sizes, we could not conduct a meaningful power analysis.

### Materials, Stimuli, and Procedure

The experiment was implemented with OpenSesame and the Expyriment back-end (Krause & Lindemann, 2014; Mathôt et al., 2012) using PyGaze for eye tracking (Dalmaijer et al., 2013). Stimuli were presented on a 27-inch monitor (1,024 × 768 pixels resolution). Using an Eyelink 1000 (SR Research Ltd., Mississauga, Ontario, Canada), participants’ pupil size and eye movements were assessed monocularly. A five-point-grid eye-tracker calibration was completed before the beginning of the task, and a single-target drift correction was performed before the start of each trial. Participants had to complete 60 trials in total, each with a timeout of 60 seconds, and were able to start the presentation of each search display themselves by pressing the space bar. Practice trials were not included in the experiment.

After the calibration, we informed participants that they would see a colorful grid of letters on the screen. Their task was to determine whether the letter grid contained an uppercase ‘F’. Subjects were instructed to press the space bar as soon as they detected the target and to press the right arrow key if they thought that the letter was absent. Overall, 224 distractor letters and either the target (‘F’) or another distractor (‘x’) were displayed in a 15×15 grid on the screen. Distractors consisted of all letters in the Latin alphabet (both upper- and lowercase), with the exception of a lowercase ‘f’ and, on target present trials, a lowercase ‘x’. The target letter appeared on approximately 50% of the trials (target presence was fully randomized). The background of the display was divided into four differently colored regions (blue, green, red, caramel/gold) that varied in shape, size, and position with each trial (Figure 1). We chose the colors so that they were roughly equal in brightness, but we did not ensure exact equiluminance.

**Figure 1.**
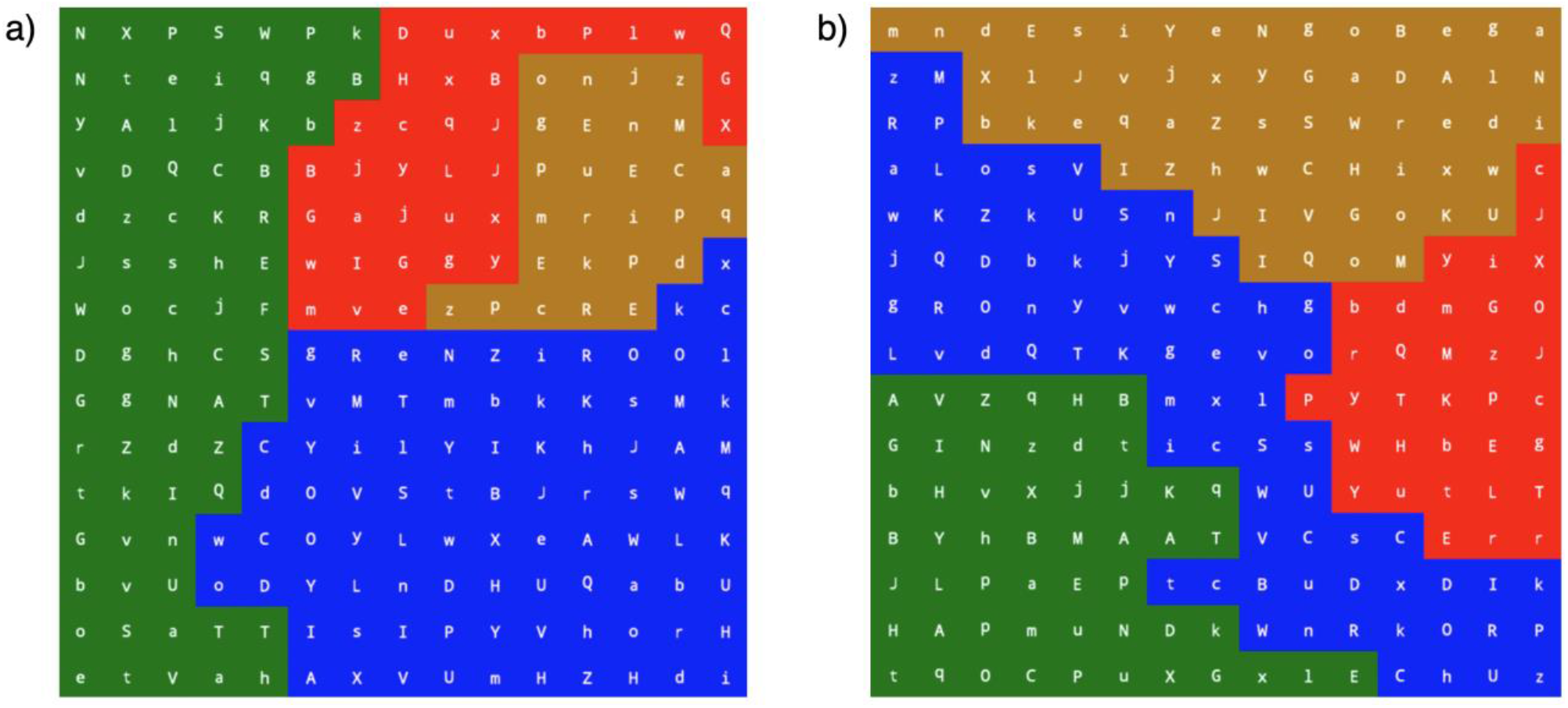
Visual-Search Trial Examples From the Experiment. *Note.* Search display with the target letter ‘F’ a) present and b) absent. All letters appeared in white, mono (Droid Sans Mono) font, 18 pixels.

Responses were followed by immediate visual feedback in the form of a red (incorrect responses) or green (correct responses) fixation dot in the center of the screen. After completion of the task, participants received final feedback on their overall response accuracy across all trials.

## Results

### Results: Behavior

Grand mean accuracy was 87.5% with a grand mean correct response time for hits of 12.4 s and 25.7 s for correct rejections.

### Preprocessing and analysis of eye-movement and pupil-size data

Blinks were reconstructed using cubic-spline interpolation (Mathôt et al., 2018). Gaze and pupil-size data were downsampled to 40 Hz. Pupil size was converted from arbitrary units as recorded by the EyeLink to millimeters of diameter using a conversion formula that we determined based on a recording of artificial pupils of known sizes. All trials and participants were included in the analysis.

We split each trial up into epochs during which gaze remained within a single region, as defined by color. A new epoch started with the start of a fixation (as determined by the EyeLink’s built-in algorithm) in a new region. To determine whether eye movements and pupil size were affected by region boundaries, we performed a ‘shuffled’ control analysis; specifically, we randomly paired the stimulus display of one trial with the data (eye movements and pupil size) from another trial, and then performed the same analyses as for the real (unshuffled) data. Whenever a pattern of results was found in the real data but not in the shuffled data, we could attribute it to an effect of regions. (The epochs in the shuffled-control analysis were slightly shorter than those in the unshuffled analysis, because participants tended to stay within regions [see below]. Therefore, we performed a third shuffled control analysis that was identical to the original shuffled analysis, except that two subsequent epochs were grouped into one, such that the epochs were longer than those in the unshuffled analysis; this did not affect the results, and the results of this analysis are therefore not further described below.)

### Results: Eye movements and Pupil Size

All Bayes Factors (BF) below result from default Bayesian paired-samples t-tests as performed in JASP 0.11.1 (JASP Team, 2020). Qualitative labels (“strong evidence”, etc.) are based on Wetzels et al. (2011).

#### Pupil size

Participants’ pupils were slightly larger before eye movements that carried gaze from one region to another than expected by chance, as shown by a comparison between the mean pupil diameter in the 1,000 ms before real (M = 4.29 mm) and shuffled region switches (Figure 2a,c; M = 4.26 mm; decisive evidence: BF_10_ = 164).

**Figure 2.**
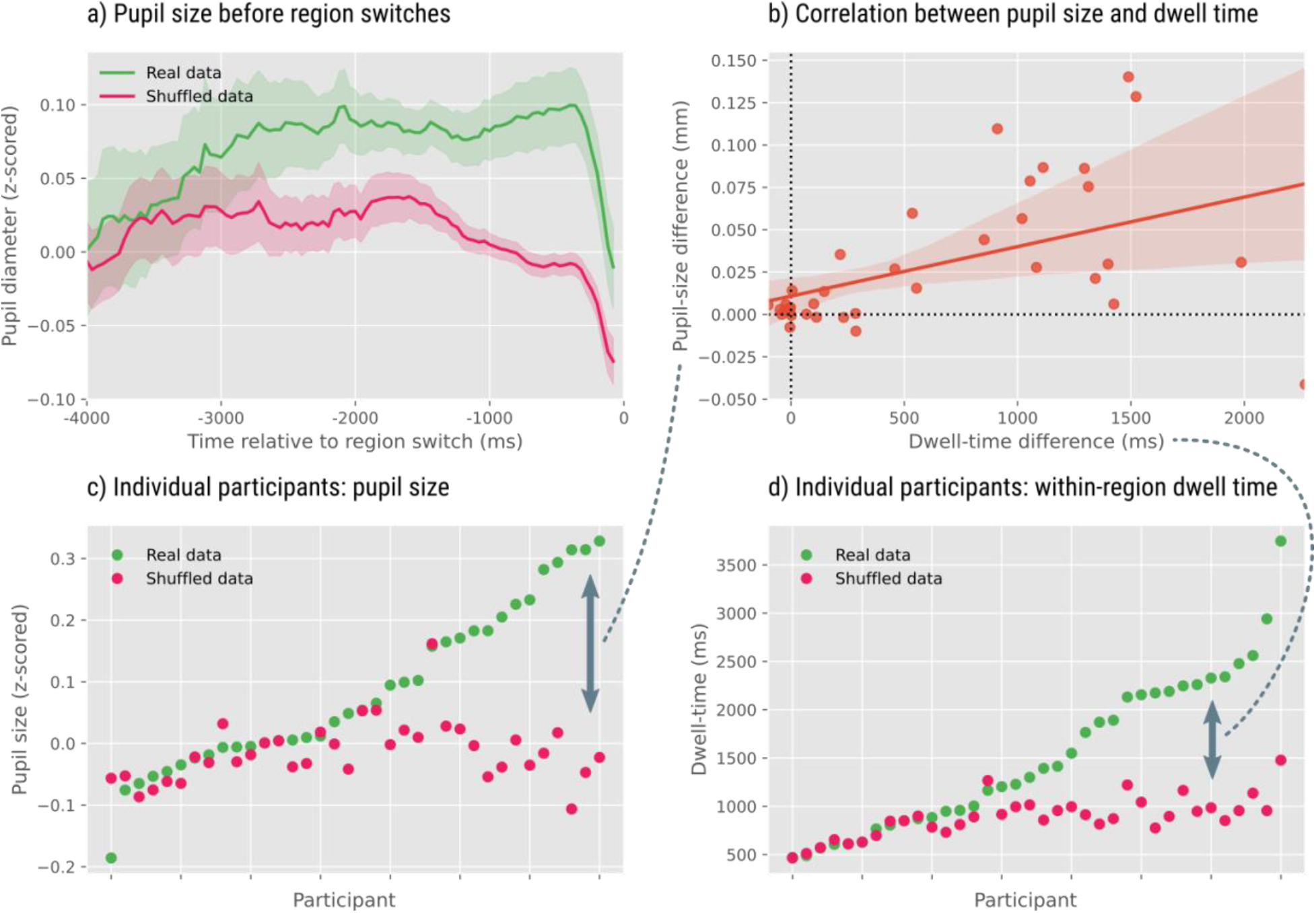
Results of Pupil Size and Eye Movement Analyses. *Note.* In all panels, green corresponds to a regular analysis of the data, whereas pink corresponds to a control analysis in which data from one trial is randomly paired with the stimulus display from another trial. Differences between green and pink indicate an effect of region. a) Pupil size (z-scored by participant) is slightly increased before eye movements that carry gaze from one region to another. The 0 point on the x-axis denotes the onset of a fixation in a new region. b) There is a positive between-subject correlation between the extent to which pupils dilate before region switches and the tendency for gaze to dwell within a region. c) Pupil size (z-scored) for all individual participants, rank-ordered by pupil size in the real analysis. d) The within-region dwell time for all individual participants, rank-ordered by dwell time in the real analysis.

#### Dwell time

Participants tended to dwell within regions for longer than expected by chance, as shown by a comparison between the within-region dwell time for the real (M = 2,435 ms) and shuffled data (Figure 2d; M = 1,420 ms; decisive evidence: BF_10_ = 9,433).

#### Correlation between dwell time and pupil size

Participants who tended to dwell within regions also showed larger pupils before switches to a new region, as shown by a correlation between the difference in dwell time between the real and shuffled analysis, and the difference in pupil diameter (as described above) between the real and shuffled analysis (Figure 2b; *r* = 0.47, BF_10_ = 10.4).

#### Size of eye movements

Eye movements that carried gaze from one region to another were slightly larger in size than would be expected by chance, as shown by a comparison between the size of region-switch eye movements for the real (M = 4.19°) and shuffled (M = 4.06°; BF_10_ = 16.3) data. To ensure that differences in eye-movement size did not explain the increase in pupil size before region switches, we conducted a Bayesian analysis of covariance (ANCOVA) with mean pupil diameter (as described above) as dependent measure, analysis type (real v shuffled) as fixed factor, subject as random factor, and eye-movement size as covariate; this analysis revealed that a model with only condition as predictor performed best, and was strongly favored over a model with only eye-movement size (very strong evidence against an eye-movement-size only model compared to an analysis-type-only model: BF_10_ = 0.012).

In summary, participants tended to dwell within regions for longer than would be expected by chance. In addition, their pupils were larger than would be expected by chance before shifts of gaze from one region to another; the effect on pupil size was numerically very small, but surprisingly robust across participants and was not explained by differences in eye-movement size. Finally, the dwell-time and pupil-size effects correlated positively with each other, suggesting that they both reflect switches from an exploitation to an exploration mode of behavior.

## Discussion

Here we show that pupils are smaller before “exploitation” eye movements that explore a single region of space than before “exploration” eye movements that carry gaze from one region to another. In addition, participants were more likely to make eye movements within regions, as compared to between regions, than would be expected by chance. These results are consistent with the adaptive-gain theory (AGT), which posits that the locus coeruleus-norepinephrine (LC-NE) system controls whether people engage in exploitation or exploration behavior, and that pupil size is a marker of LC activity. Previous studies have tested the AGT with complex, game-like tasks. Crucially, here we show that the AGT also applies to a very basic form of behavior: eye movements.

### Central Versus Peripheral Focus

Although our findings are correlational, the results of the present study support the AGT’s notion that eye movements and changes in pupil size are linked to behavioral modes. During the search task, participants’ bias to make eye movements within a region is suggestive of exploitative behavior and was accompanied by smaller pupils. Notably, we observed that pupils dilated just before subjects made an eye movement to another region, indicating a shift from exploitative to exploratory behavior. These adjustments in pupil size might represent a form of “sensory tuning” (Mathôt, 2020); specifically, a shift in focus to either central or peripheral parts of our visual field, accompanied by a small change in pupil diameter, could slightly improve visual input for subsequent sensory processing. The pupil-size changes that we observed here, during a simple visual-search task in the laboratory, were very small; however, similar adjustments in pupil diameter could be larger, and thus have bigger effects, in real-life scenarios. Taken together, small adaptations in pupil size that coincide with switches in behavioral modes (exploitative versus exploratory) could serve a function by optimizing vision for the given situation.

### Object-Based Attention

The tendency to make eye movements within regions, as opposed to between regions, is reminiscent of object-based attention. The notion of object-based attention posits that when we attend to one part of an object, visual attention spreads within the boundaries of that object. A similar within-object preference also holds for eye movements. For instance, Theeuwes et al. (2010) employed a classic cueing paradigm (Egly et al., 1994) in which two adjacent rectangles were presented either horizontally or vertically on the screen. While participants fixated on a central dot, the end of one of the four rectangles was cued with a salient, task-irrelevant cue that captured attention. Immediately after, three distractors and one target letter appeared, one letter in each of the rectangles’ corners. Letters were too small to be identified through a covert shift of attention alone, and participants therefore had to make eye movements towards the letters to identify them. As to be expected, participants generally first looked at the cued location. Crucially, however, participants were more likely to make the second eye movement to the letter within the same rectangle than to an equidistant letter in the other rectangle. This finding is reminiscent of the results of the current study, and we therefore propose that object-based attention and exploitation behavior are different ways to look at the same fundamental behavior; that is, attention preferentially spreads within an object because we tend to ‘exploit’ this object before switching to another object. An important avenue for future research will be to bridge the gap between the literature on visual-attention and the literature on arousal-related pupil dilation, for example by testing whether the predictions of the AGT extend to the kind of simple cuing paradigms that are generally used to study object-based attention (for a recent study that addressed this topic in a different way, see O’Bryan & Scolari, 2021).

### Conclusion

In conclusion, eye movements that carry gaze from one region (defined by color) to another region are accompanied by a slight dilation of the pupil. This suggests that the distinction between exploitation and exploration, as posited by the AGT, extends from complex game-like tasks to simple visual-search tasks. We further propose that the concepts of exploitation and exploration are closely related to object-based attention.

